# Novel Approaches to Track Neurodegeneration in Murine Models of Alzheimer’s Disease Reveal Previously Unknown Aspects of Extracellular Aggregate Deposition

**DOI:** 10.1101/2025.04.04.647232

**Authors:** Daisy Gallardo, Oswald Steward

## Abstract

This paper describes a novel transgenic-based platform to track degeneration of specific populations of neurons in 5xFAD mice, a murine model of Alzheimer’s disease. We created a new double transgenic model by crossing 5xFAD mice with Rosa^tdT^ reporter mice. 5xFAD^+/−^/Rosa^tdT^ mice received intra-spinal cord injections of AAV-retrograde (rg)/Cre at 2-3 months of age to permanently label corticospinal neurons (CSNs). Brains and spinal cords were retrieved 2-3 weeks post-injection or at 12-14 months of age. Immunohistochemical studies of transgene expression throughout the brain and spinal cord, using an antibody selective for hAPP, revealed age-dependent accumulation of hAPP in extracellular aggregates in regions containing hAPP expressing neuronal cell bodies and in regions containing axons and synaptic terminals from hAPP expressing neurons. Permanent labeling of CSNs with tdT confirmed extensive loss of CSNs in old mice. Surprisingly, we discovered that tdT expressed by CSNs accumulated in extracellular aggregates that persisted after the neurons that expressed tdT degenerated. Extracellular aggregates of tdT also contained hAPP and co-localized with other markers of AD pathology. Overall, deposition of hAPP in extracellular aggregates in areas containing axons and synaptic terminals from hAPP expressing neurons is a prominent feature of AD pathophysiology in 5xFAD mice. In addition, accumulation of hAPP and reporter proteins in extracellular aggregates provides a secondary measure to track neurodegeneration of identified populations of neurons in these mice.

**Highlights:**

– Characterization of a new double transgenic strain allowing Cre-dependent labeling of populations of neurons that degenerate in 5xFAD mice.
– Selective labeling of layer V corticospinal neurons (CSNs) via retrograde transduction with AAV-rg allows quantification of previously un-recognized aspects of age-dependent CSN degeneration.
– Age-dependent deposition of extracellular hAPP by axons and synaptic terminals revealed by immunohistochemistry for mutant human APP in 5xFAD mice
– Petal-shaped clusters of hAPP originate mainly due to axonal degeneration and fragmentation.
– Surprisingly, tdTomato expressed by neurons that degenerate, persists in extracellular deposits that co-localize with extracellular deposits of hAPP.

## 1. Introduction

Alzheimer’s Disease (AD) is one of the most prevalent neurodegenerative diseases with an estimated 6.9 million Americans aged 65 and older currently affected (2024 Alzheimer’s disease facts and figures). Early in disease onset, AD is characterized by the accumulation of extracellular amyloid beta (Aß) plaques and intra-neuronal neurofibrillary tangles (Terry, 1963). In late-stage AD, neurodegeneration is a primary pathological feature. Neurodegeneration includes dendritic atrophy, axonal destabilization and finally neuron death (Terry et al., 1991; Thal et al., 2002). Extracellular aggregated Aß deposits are commonly surrounded by degenerating neuronal processes, and these together are termed neuritic plaques. Some of these pathological features are recapitulated in animal models of Alzheimer’s disease (Drummond and Wisniewski, 2016). Fully characterizing the progression of pathology in these models is crucial for the understanding of disease mechanisms and development of effective treatments.

A commonly used murine model of Alzheimer’s disease is the 5xFAD transgenic strain, which expresses two mutant human genes [amyloid precursor protein (APP) and presenilin 1 (PSEN1)] with 5 familial Alzheimer’s mutations (3 in APP; 2 in PSEN1). During the creation of 5xFAD mice, the two transgenes are inserted into a single locus and are driven by the neuron-specific elements of the mouse Thy-1 promoter (Oakley et al., 2006). Using an antibody that recognizes both mouse and human proteins, total levels of APP protein expression in brain (mouse and human) have been found to be 1.8 times than in non-transgenic control mice (Sadleir et al., 2018).

As they age, 5xFAD mice have been reported to develop high Aβ_42_ levels, amyloid plaques, and exhibit decreases in synaptic and neuronal markers in the brain (Oakley et al., 2006). Of note, 5xFAD mice are one of the only AD transgenic strains that exhibit actual neurodegeneration; neurodegeneration has been reported to occur in cortical layer V and the subiculum. Cortical layer V and the subiculum are areas where the transgenes are highly expressed based on them being driven by Thy1 (Moechars et al., 1996; Oakley et al., 2006) and are areas with high amyloid plaque burden (Oakley et al., 2006; Eimer and Vassar, 2013).

Despite extensive use of 5xFAD mice, there have been limited studies of the overall pattern of transgene expression in different neuron types throughout the brain and spinal cord. Additionally, there have been limited studies on whether different neuron types that express the transgene exhibit age-dependent transgene expression and/or degenerative changes. One goal of this study is to explore this question by immunohistochemistry using a commercially available antibody (Anti-APP, clone 1D1, Millipore Cat# MABN2287, RRID:AB_3665093), which we show here selectively recognizes the mutant human APP (hAPP) in 5xFAD mice and not mouse APP (mAPP). The 1D1 antibody has been reported to bind to the N-terminus of hAPP, amino acids 40-64 of its unmodified state, detecting both the membrane bound and soluble hAPP (Hofling et al., 2016). In the Hofling et al., 2016 report, two monoclonal antibodies against hAPP were created, clone 1D1 and clone 7H6. Prior to the 1D1 and 7H6 antibodies, there were no available antibodies that selectively detected human APP, but not mAPP, in brains of APP-transgenic mice. We chose to use the 1D1 antibody which exhibits identical immunoreactivity to the 7H6 antibody (Hofling et al., 2016). Another widely used antibody, 6E10 has previously been proposed to selectively identify hAPP and not endogenous mAPP; however, this antibody has been found to react with mAPP in non-reducing conditions by western blot analysis (Hofling et al., 2016). In addition to APP, 6E10 recognizes Aβ in immunocytochemical preparations, enzyme linked immunosorbent assays (ELISA) and western blots from APP-transgenic mouse tissue, while 1D1 does not (Herzig et al., 2004; Zheng et al., 2012; Hofling et al., 2016).

The second goal of this study was to develop a technique that selectively and permanently labels neurons that degenerate in 5xFAD mice, specifically targeting layer V pyramidal neurons in the sensorimotor cortex. The purpose of the technique is to allow quantitative analyses of the time course and characteristics of neurodegeneration. Previous reports of neurodegeneration in 5xFAD mice have used routine neurohistological stains such as Cresyl violet to document decreases in numbers of large cells in layer V of the somatosensory cortex. To our knowledge, there have been no studies using selective markers to label specific populations of neurons. For example, although it is reasonable to speculate that some of the layer V neurons that degenerate are the neurons that give rise to the corticospinal tract, this has not been established. For this purpose, we developed a new strain of double-transgenic 5xFAD mice that allows selective and permanent retrograde labeling of layer V cortical neurons and other neuron types in the brain that project to the spinal cord. Full labeling of dendrites and axons allows tracking of features of neurodegeneration across age. This model has allowed specific identification of age dependent degeneration of layer V corticospinal neurons (CSNs) as well as other neuron types that have not previously been reported to degenerate in 5xFAD^+/−^/Rosa^tdT^ mice. We also document early degenerative changes in axons and dendrites and discovered that extracellular tdT particles accumulate as neurons degenerate, providing a secondary tool to track neurodegeneration from an identified population of neurons.

## 2. Methods

### 2.1 Experimental Animals

All procedures involving animals were approved by the Institutional Animal Care and use Committee (IACUC) at the University of California, Irvine (UCI) in accordance with the National Institute of Health (NIH) guidelines. Mice had ad lib access to food and water and were housed at 25°C with a 12h light/dark cycle. To produce the transgenic mice for these studies, 5xFAD (B6SJL-Tg (APPSwFlLon, PSEN1*M146L*L286V) 6799Vas/Mmjax; RRID:MMRRC_034840-JAX) mice were bred with Rosa^tdTomato(tdT)^ mice resulting in 5xFAD/Rosa^tdT^ and Rosa^tdT^ littermate controls. 5xFAD mice were obtained from Jackson Laboratory (Stock No: 34840-JAX). The Rosa^tdT^ mice were originally obtained from Jackson Laboratory (Stock No: 007905) and maintained in our breeding colony over multiple generations before being crossed with 5xFAD mice.

In 5xFAD mice, the origin of transgene inheritance has been reported to influence amyloid plaque burden. 5xFAD mice with maternally inherited transgenes were shown to have a slower progression of amyloid plaque formation (Sasmita et al., 2024). The report highlighted the importance of reporting the origin of transgenes. Below, in Table 1, we have provided details on the parental origin of transgenes in the mice used for these studies.

**Table 1.** Mice used for these studies.

| *Injected with AAV-rg/Cre at 2-3 months of age* |  | **Months of age at collection** |  |  |  |
| --- | --- | --- | --- | --- | --- |
| **Genotype** | **Parental origin of transgenes** | **2-4** | **6-8** | **10-12** | **14-15** |
| 5xFAD<sup>+/-</sup>/Rosa<sup>tdT</sup> | Both | 5M/1F | 1M | 1M/6F | 2M/2F |
|  | Maternal | - | - | - | - |
|  | Paternal | 4M/1F | - | 2M | 1M/4F |
| Rosa<sup>tdT</sup> | No transgenes | 4M/16F | 1F | 1M | 4F |
| *Injected with AAV-rg/Cre at various ages* |  | **Months of age at collection** |  |  |  |
| **Genotype** | **Parental origin of transgenes** | **4-5** | **6-8** | **9-12** | **13-29** |
| 5xFAD<sup>+/-</sup>/Rosa<sup>tdT</sup> | Both | 2M/2F | 15M/9F | 6M/7F | - |
|  | Maternal | - | - | - | - |
|  | Paternal | - | - | - | - |
| Rosa<sup>tdT</sup> | No transgenes | 1M | 7M/4F | 1M/1F | 2M |

**Table 2.** Primary antibody list.

| **Antibody** | **Host** | **Vendor** | **Catalog #** | **RRID** | **Dilution** |
| --- | --- | --- | --- | --- | --- |
| Human APP, clone 1D1 | Rat | Millipore Sigma | MABN2287 | RRID:AB_3665093 | 1:250 |
| RFP | Rabbit | Rockland | 600-401-379 | RRID:AB_2209751 | 1:1000 |
| LAMP1 | Rabbit | Abcam | ab24170 | RRID:AB_775978 | 1:300 |
| NeuN, clone A60 | Mouse | Millipore Sigma | MAB377 | RRID:AB_2298772 | 1:250 |
| Human Aβ | Mouse | Tecan (IBL) | 10323 | RRID:AB_10707424 | 1:1000 |

### 2.2 Surgical procedures

At 2-3 months of age, mice received intra-spinal cord injections of AAV-retrograde(rg)/Cre (Addgene viral prep #24593-AAVrg) at cervical level 5 (C5). Mice were anesthetized with continuous Isoflurane (2-2.5%) administration, their eyes were swabbed with Vaseline to prevent drying, and scalp was shaved and cleaned with Betadine®. A laminectomy was performed at C5 to expose the spinal cord and AAV-rg/Cre was injected 0.5 mm left of the midline and 0.5 mm deep (3E9 viral genome copies (GC) per mouse) using a Hamilton microsyringe fitted with a pulled glass micropipette. The needle was kept in place for 1 minute after completing the injection. Following injection, the muscle layers were sutured, and the skin was closed by wound clips.

### 2.3 Post-operative care

After surgery, mice were immediately placed in cages with Alpha-Dri bedding with half of the cage placed on a circulating water warming pad set to 37°C. Partial placement of cage on top of warming pads allows mice to remove themselves from the heat source, if desired. Mice were kept on the warming pad for 3 days post-surgery and received lactated Ringer’s solution once a day (1ml/20g, subcutaneously), enrofloxacin once a day (Baytril® 2.5mg/kg, subcutaneously) and buprenorphine twice a day (0.01mg/kg, subcutaneously). Mice were monitored twice daily for 3 days post-injury for general health and signs of distress. If bladders of mice were full, indicating difficulty to urinate due to surgery, the bladders were manually expressed twice daily. Bladder expression was required for a small number of animals and for a maximum of 3 days post-surgery. After 3 days, the cages were removed from the warming pads and were transferred back to the mouse housing room set at 25°C. Once in the housing room, mice were routinely monitored by vivarium staff and research staff.

### 2.4 Tissue collection

All mice were euthanized using Fatal Plus®, followed by transcardiac perfusion with 4% paraformaldehyde (PFA). Brains and spinal cords were removed intact and imaged under a fluorescent microscope to visualize tdT fluorescence from the site of injection to the cortex. Brains and spinal cords were kept in 4% PFA for at least 24 hours. The tissue was then submerged in 27% sucrose for 24 hours for cryoprotection and frozen in Tissue-Tek O.C.T. embedding medium. Brains were then sectioned at 30µm in coronal or horizontal planes (*see Results*), using a cryostat. Sections were stored free-floating in phosphate-buffered saline (1xPBS, 20mM, pH7.4) with 0.1% sodium azide.

### 2.5 Immunohistochemistry (IHC) and immunofluorescence (IF)

Free-floating brain sections at 480 µm intervals were used for IHC and IF. Human APP transgene expression was detected by immunostaining for Anti-APP, clone 1D1 (Millipore Cat# MABN2287, RRID:AB_3665093). For 1D1 IF, sections were washed in Tris Buffered Saline (1xTBS, 100mM Tris. pH 7.4 and 150 mM NaCl) 2 times for 5-10 minutes each time. The sections were then incubated in methanol with 3% Hydrogen peroxide for 15 minutes and washed in 1xTBS 3 times for 5-10 minutes each. After washing, sections were incubated for 1-2 hours at room temperature in a 1xTBS, 5% Normal Donkey Serum (NDS), 0.3% Triton X-100 blocking solution. Sections were then incubated in a 1:250 dilution of Rat Anti-1D1 primary antibody (Millipore Cat# MABN2287, RRID:AB_3665093) with 1xTBS, 5% NDS and 0.3% Triton X-100 overnight at room temperature. The following day, sections were washed in 1xTBS 3 times for 5-10 minutes each and then incubated in a 1:250 dilution of Biotin-SP (long spacer) AffiniPure™ Donkey Anti-Rat IgG (H+L) (Cat# 712-065-153, RRID: AB_2315779, Jackson ImmunoResearch) for 2 hours at room temperature. Sections were then washed in 1xTBS 3 times for 5-10 minutes each and then incubated in a 1:100 dilution of VECTASTAIN® Elite® ABC (Avidin/Biotin) - Horseradish peroxidase (HRP) (Standard) (PK-6100, RRID: AB_2336819) with 1xTBS for 1-2 hours. After incubation, sections were washed in 1xTBS 3 times for 5-10 minutes each and then incubated in a 1:250 dilution of Fluorescein isothiocyanate (FITC) with 0.1M borate buffer (pH 9) for 25 minutes. Sections were washed twice in 1xTBS and mounted on 0.5% gelatin subbed, double frosted, VWR® microscope slides. For NeuN IF, sections were washed in Tris Buffered Saline (1xTBS, 100mM Tris. pH 7.4 and 150 mM NaCl) 2 times for 5-10 minutes each time. The sections were then incubated in methanol with 3% Hydrogen peroxide for 15 minutes and washed in 1xTBS 3 times for 5-10 minutes each. After washing, sections were incubated for 1-2 hours at room temperature in a 1xTBS, 5% NDS, 0.3% Triton X-100 blocking solution. Sections were then incubated in a 1:250 dilution of Anti-NeuN Antibody, clone A60 (Millipore Cat# MAB377, RRID:AB_2298772) in 1xTBS, 5% NDS, 0.3% Triton X-100 solution overnight at room temperature. The following day, sections were washed in 1xTBS 3 times for 5-10 minutes each and then incubated in a 1:250 dilution of Donkey anti-Rat HRP conjugated secondary antibody with 1xTBS, 5% NDS, 0.3% Triton X-100 for 2 hours at room temperature. Sections were washed in 1xTBS 3 times for 5-10 minutes each and then incubated in a 1:250 dilution of Donkey anti-mouse HRP conjugated secondary antibody with 1xTBS, 5% NDS, 0.3% Triton X-100 for 2 hours at room temperature. Then, sections were washed 3 times in 1xTBS and incubated in a 1:250 dilution of Fluorescein isothiocyanate (FITC) with 0.1M borate buffer (pH 9) for 25 minutes. Sections were washed twice in 1xTBS and mounted on 0.5% gelatin subbed, double frosted, VWR® microscope slides.

To detect tdTomato by IF, sections were immunostained for Red Fluorescent Protein (RFP). Steps prior to incubation in primary antibody were as for 1D1 (*above*). Sections were then incubated in a 1:1000 dilution of Rabbit anti-RFP primary antibody (Rockland Cat# 600-401-379, RRID:AB_2209751) with 1xTBS, 5% NDS and 0.3% Triton X-100 overnight at room temperature. The next day, sections were washed in 1xTBS 3 times for 5-10 minutes each and then incubated in a 1:500 dilution of Donkey anti-Rabbit 555 with 5% NDS and 0.3% Triton X-100 for 2 hours at room temperature, washed twice in 1xTBS and mounted on 0.5% gelatin subbed, double frosted, VWR® microscope slides.

For the RFP and 1D1 co-IF, the steps and reagents remained the same for each, but the steps were combined. The Aβ IF was done by first washing sections in 1xTBS 2 times for 5-10 minutes each, then sections were incubated in 1.75 mL clear microtubes filled with citrate buffer (10mM, pH 6) and tubes were placed in a 100 degrees Celsius water bath for 5 minutes for antigen retrieval and then in a cool water bath for 10 minutes. Sections were washed in 1xTBS 3 times for 5-10 minutes each, incubated in 88% formic acid (Cat# A119P-500, Fisher Scientific) for 7 minutes, then washed again in 1xTBS 3 times for 5-10 minutes each. Sections were then incubated for 1-2 hours at room temperature in a 1xTBS, 5% NDS, 0.3% Triton X-100 solution followed by incubation in a 1:1000 dilution of Mouse anti-Amyloid-ß (N) (82E1) Aβ Anti-Human Mouse IgG MoAb (Tecan (IBL) Cat# 10323, RRID:AB_10707424) with 1xTBS, 5% NDS, 0.3% Triton X-100 overnight at room temperature. The following day, sections were washed in 1xTBS 3 times for 5-10 minutes each and then incubated in a 1:250 dilution of Donkey anti-Mouse 488 with 1xTBS, 5% NDS, 0.3% Triton X-100 for 2 hours at room temperature. Lastly, sections were washed twice in 1xTBS and mounted on 0.5% gelatin subbed, double frosted, VWR® microscope slides. Some sections were stained with Hoechst in 1xPBS (1µg/ml) to label cell nuclei. The LAMP1 IF was done by incubating sections in a 1xTBS, 5% NDS, 0.3% Triton X-100 solution for 1-2 hours at room temperature, followed by an overnight incubation in a 1:300 dilution of Rat anti-LAMP1 (Abcam Cat# ab24170, RRID:AB_775978) in 1xTBS, 5% NDS, 0.3% Triton X-100. The following day, sections were washed in 1xTBS 3 times for 5-10 minutes each and then incubated in a 1:250 dilution of Donkey anti-Rat-biotinylated conjugated secondary antibody with 1xTBS, 5% NDS, 0.3% Triton X-100 for 2 hours at room temperature. Then, sections were washed 3 times in 1xTBS and incubated in a 1:100 dilution of VECTASTAIN® Elite® ABC-HRP, Peroxidase (Standard) (PK-6100, RRID: AB_2336819) with 1xTBS for 1-2 hours. After incubation, sections were washed in 1xTBS 3 times for 5-10 minutes each and then incubated in a 1:250 dilution of Fluorescein isothiocyanate (FITC) with 0.1M borate buffer (pH 9) for 25 minutes. Lastly, sections were washed in 1xTBS twice for 5-10 minutes each and mounted on 0.5% gelatin subbed, double frosted, VWR® microscope slides.

### 2.6 Quantification of corticospinal neurons (CSNs)

For quantification of tdT-labeled neurons, coronal sections at 240 µm intervals were mounted without immunostaining onto 0.1% gelatin subbed slides. Sections were not immunostained because with IHC, robust labeling of dendrites can obscure tdT-labeled CSN cell bodies. TdT-positive CSNs in layer V of the cortex were manually counted in each section from the medial border of M1 to the lateral border of S1 using a fluorescent Olympus BX53 microscope at 20x magnitude. The population of TdT-labeled CSNs in the dorso-lateral cortex posterior to bregma that are separated from the main sensorimotor cortex were not counted. Counts were taken in all sections in the series that contained tdT-labeled CSNs, which typically was from 3 mm anterior to bregma to approximately 2.5 mm posterior to bregma. Importantly, only tissue from 5xFAD^+/−^/Rosa^tdT^ and Rosa^tdT^ mice injected between 2-3 months old was used for quantifications.

### 2.7 Quantification of extracellular accumulations of tdT (debris)

The right cortex of a coronal section at approximately 0.74mm–0.62mm anterior to bregma was selected and imaged with a fluorescent Olympus BX53 microscope at 20x magnification. The quantifications were on the right cortex because the unilateral AAV-rg/Cre injection was on the left side of the spinal cord. Multiple images were taken at 10x magnification to span the area containing labeled CSNs, images were combined into a montage and imported to Fiji ImageJ. The scale was set to pixels and images were converted to 8-bit format. Then, the threshold was set to 0.36–0.38% with a dark background. After thresholding, the watershed separation command was used to accurately track the clumps of debris. If any neurons were selected throughout the process (identified by their morphology and location in layer V), they were removed. Lastly, the particle analysis plug-in was used with the following settings: Size (pixel^2^) = 100–1000, circularity = 0.30–1.00.

## 3. Results

### 3.1 Pattern of expression of AD transgenes in brain and spinal cord of young 5xFAD/Rosa^tdT^ mice

For this study, we created a new transgenic strain by crossing 5xFAD mice (34840-JAX) with Rosa^tdTomato^ reporter mice. These mice allow for selective and permanent retrograde labeling of corticospinal neurons (CSNs) which are amongst the neurons in cortical layer V that degenerate in 5xFAD mice. Since these mice have a different genetic background than standard 5xFAD mice, it was important to determine the overall pattern of expression of the 5xFAD transgenes. Also, to our knowledge, there have been no comprehensive studies of the overall pattern of expression of hAPP in different neuronal cell types in the parental 5xFAD transgenic strain. The mutant hAPP and hPS1 genes in 5xFAD mice are driven by a modified mouse Thy-1 promoter, but the exact pattern of Thy-1 driven expression can vary. A previous study described patterns of labeling in cortex, hippocampus and spinal cord using an unpurified polyclonal antiserum against human APP, but specificity for hAPP was not documented (Jawhar et al., 2012). A more recent study illustrated examples of APP expression in cortex and hippocampus, but only these brain regions were shown and, again, the antibody used was not specific to hAPP.

To determine the pattern of expression in our transgenic strains, we immunostained coronal brain sections from young (2-month-old) 5xFAD^+/−^/Rosa^tdT^ and Rosa^tdT^ mice using an antibody for hAPP (Clone 1D1, Millipore Cat# MABN2287, RRID:AB_3665093), with FITC as the reporter. To test selectivity of this antibody for hAPP, we also immunostained sections from control mice lacking the 5xFAD transgene (Rosa^tdT^ mice).

Immunofluorescence for hAPP was completely absent in control Rosa^tdT^ mice (**Fig. 1A**) whereas there was robust labeling of multiple populations of neurons in 5xFAD^+/−^/Rosa^tdT^ mice (**Fig. 1B**). This documents that with our methods of immunostaining, the 1D1 antibody specifically recognizes hAPP and not mouse APP. To our knowledge, the monoclonal 1D1 antibody has not previously been used to document expression of hAPP in 5xFAD mice, so the following describes the overall pattern of expression in some detail. For this, we first describe the pattern of labeling in young (2–3-month-old) mice and the change in the pattern of labeling as the mice age.

**Figure 1.**
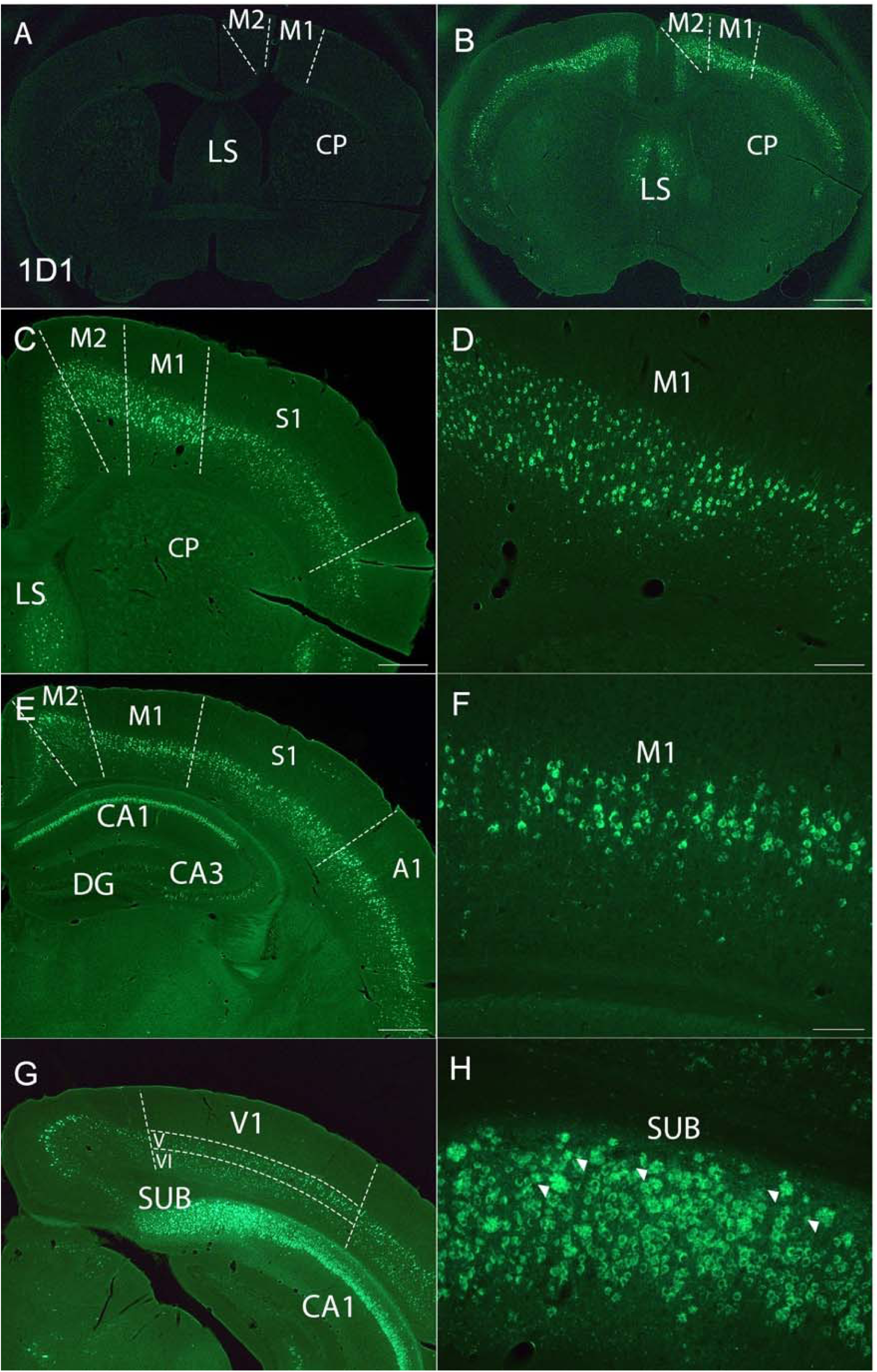
Immunofluorescence for hAPP in the cortex, hippocampus and entorhinal cortex of 2-month-old 5xFAD^+/−^/Rosa^tdT^ and Rosa^tdT^ mice. **(A)** Rosa^tdT^ mouse stained with 1D1 antibody against hAPP. Absence of labeling indicates specificity for hAPP, indicating no cross-reactivity with mouse APP. No immunofluorescence in primary (M1) and secondary (M2) motor cortex, lateral septum (LS) and caudate putamen (CP). **(B)** Pattern of immunofluorescence for hAPP in 5xFAD^+/−^/Rosa^tdT^ mouse in section approximately 1mm anterior to bregma at the level of the primary motor cortex. **(C)** Higher magnification view of labeling of neurons in M1, M2, primary somatosensory cortex (S1) and LS. No apparent labeling in the dentate gyrus (DG). **(D)** Higher magnification view of M1. **(E)** Pattern of immunofluorescence in cortex and hippocampus approximately 2mm caudal to bregma. Note lower levels of labeling in CA3 vs. CA1 region of the hippocampus. **(F)** Higher magnification view of M1 cortex shown in E. **(G)** Pattern of immunofluorescence in layer V/VI of the visual cortex, CA1 and subiculum (SUB), approximately 3.5 mm caudal to bregma. **(H)** Higher magnification view of subiculum and presence of 1D1 immunofluorescent rosette shaped clumps (arrowheads). Scale bar: 1mm in **A & B**, 500 µm in **C, E, G,** 200µm in **D, F & H**.

As expected, based on previous studies, neurons in layer V of the primary (M1) and secondary (M2) motor area and primary somatosensory (S1) area exhibited high levels of 1D1 immunofluorescence (**Fig. 1B-F**). Importantly, these are the neurons that have been reported to degenerate in 5xFAD mice (Oakley et al., 2006). Neurons in layers IV and VI were also labeled although fluorescence was less pronounced than in layer V.

In sections taken near the caudal boundary of the sensorimotor cortex (approximately 1.7 mm posterior to bregma), there were fewer 1D1-positive neurons in layer V (**Fig. 1E&F**) than in the more rostral M1 region (**Fig. 1D**). Labeling was more prominent in layer V and VI of the S1 area and auditory (A1) region in the lateral cortex (**Fig. 1E**). In more caudal sections (approximately 3.5mm posterior to bregma) where the visual cortex was prominent, labeling was also present in layers V and VI (**Fig. 1G).**

A high level of immunofluorescence for hAPP was also observed in the pyramidal layer of the CA1 region of the hippocampus (**Fig. 1E**) and in more caudal sections, in the subiculum (**Fig. 1G&H**). Levels of labeling in CA2-3 were lower than CA1 but still detectable (**Fig. 1E**). Of note, in addition to labeled neuronal cell bodies in the subiculum, there were clusters of 1D1-positive particles (**Fig. 1H**). These clusters had a distinctly different appearance than the 1D1-positive neuronal cell bodies in the cell layer of the subiculum in that the particles appeared to be extracellular (more on extracellular 1D1-positive particles below). Of note, subicular neurons have been reported to degenerate in 5xFAD mice (Oakley et al., 2006) whereas minimal, if any, neurodegeneration has been reported in CA1 and CA3.

In the brains of patients with AD, the entorhinal cortex is one of the first regions to show signs of neurofibrillary tangles and neurodegeneration (Braak and Braak, 1991, Gomez-Isla et al., 1996). To our knowledge, the pattern of expression of 5xFAD transgenes has not been characterized in the entorhinal cortex or posterior cortical regions with antibodies specific to human APP. Accordingly, we sectioned brains in the horizontal plane from 3-month-old 5xFAD^+/−^/Rosa^tdT^ mice to better visualize the entorhinal cortex and other posterior cortical areas and immunostained these for hAPP with the 1D1 antibody. Figure 2 illustrates the pattern of labeling in sections through the dorsal EC and in a more ventral section where the lateral EC is present. 1D1-positive neurons with pyramidal morphology were present in layers V and VI of the entorhinal cortex (**Fig. 2A&B**). Notably, there is minimal hAPP fluorescence in stellate neurons in layer II of the entorhinal cortex, which are the cells of origin of perforant path projections to the dentate gyrus and distal dendrites of CA3 (Steward and Scoville, 1976). The lack of labeling for hAPP in layer II is noteworthy because in humans, neurons in layer II of the entorhinal cortex are the most susceptible to early neurodegeneration in AD patients (Gomez-Isla et al., 1996). In addition, there was little labeling of the pyramidal neurons in layer III, which project to the most distal dendrites of neurons in CA1-subiculum (Steward and Scoville, 1976). Similarly, there was essentially no 1D1 labeling in the parasubiculum and presubiculum. There was a high level of labeling for 1D1 in neurons in the subiculum and accumulations of 1D1-positive extra-cellular clusters of 1D1 particles in the *stratum oriens* were even more apparent than in the dorsal subiculum visible in coronal sections (**Fig. 2A&B**).

**Figure 2.**
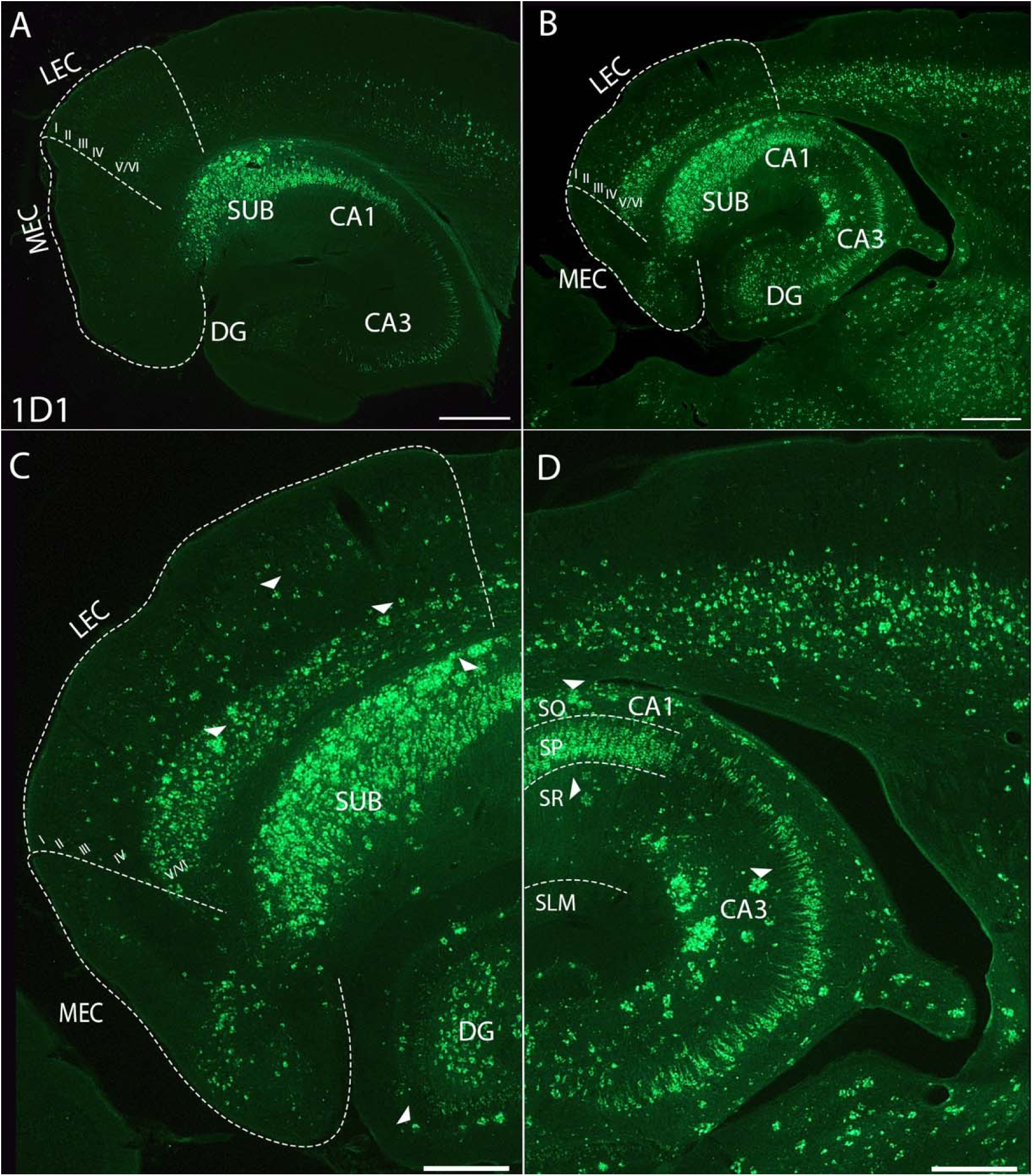
Sections on a horizontal plane showing hAPP fluorescence. **(A & B)** Layers of the MEC and LEC in a 3-month-old 5xFAD^+/−^/Rosa^tdT^ mouse. Immunofluorescence for hAPP also seen throughout SUB, CA1, CA2, CA3 and DG. **(C & D)** In 5xFAD^+/−^/Rosa^tdT^ mice of 10 months of age, abundant presumably intraneuronal immunofluorescence for hAPP seen throughout SUB and CA1, with lower amounts seen in CA3 and DG. 1D1 positive clusters scattered throughout the SUB, MEC, LEC DG **(arrowheads, C)**, *stratum oriens* (SO) and *stratum radiatum (*SR) of CA1 and in CA3 **(arrowheads, D)**. **(A & C)** Sections at about −3.0 mm from Bregma in dorsoventral axis. **(B & D)** Sections at about −3.96 mm from Bregma in dorsoventral axis. Scale bar: 500 µm in **A & B**, 300µm in **C & D.**

In subcortical regions of 2–3-month-old mice, there was moderate 1D1 labeling in the lateral septum (**Fig. 1B** and **Fig. 3A & A1, LS**), and in neurons of the basolateral amygdala (BLA, **Fig. 3B & B1).** The fluorescent labeling in the lateral septum, which receives glutamatergic input from the hippocampus (Khakpai et al., 2013), did not appear to be inside of neuronal cell bodies but rather appears to be extracellular. This is distinct from the fluorescent labeling seen in neuronal somata of cortical and hippocampal neurons (**Fig. 1E & F**) and neurons in the basolateral amygdala (BLA, **Fig. 3B & B1).**

**Figure 3.**
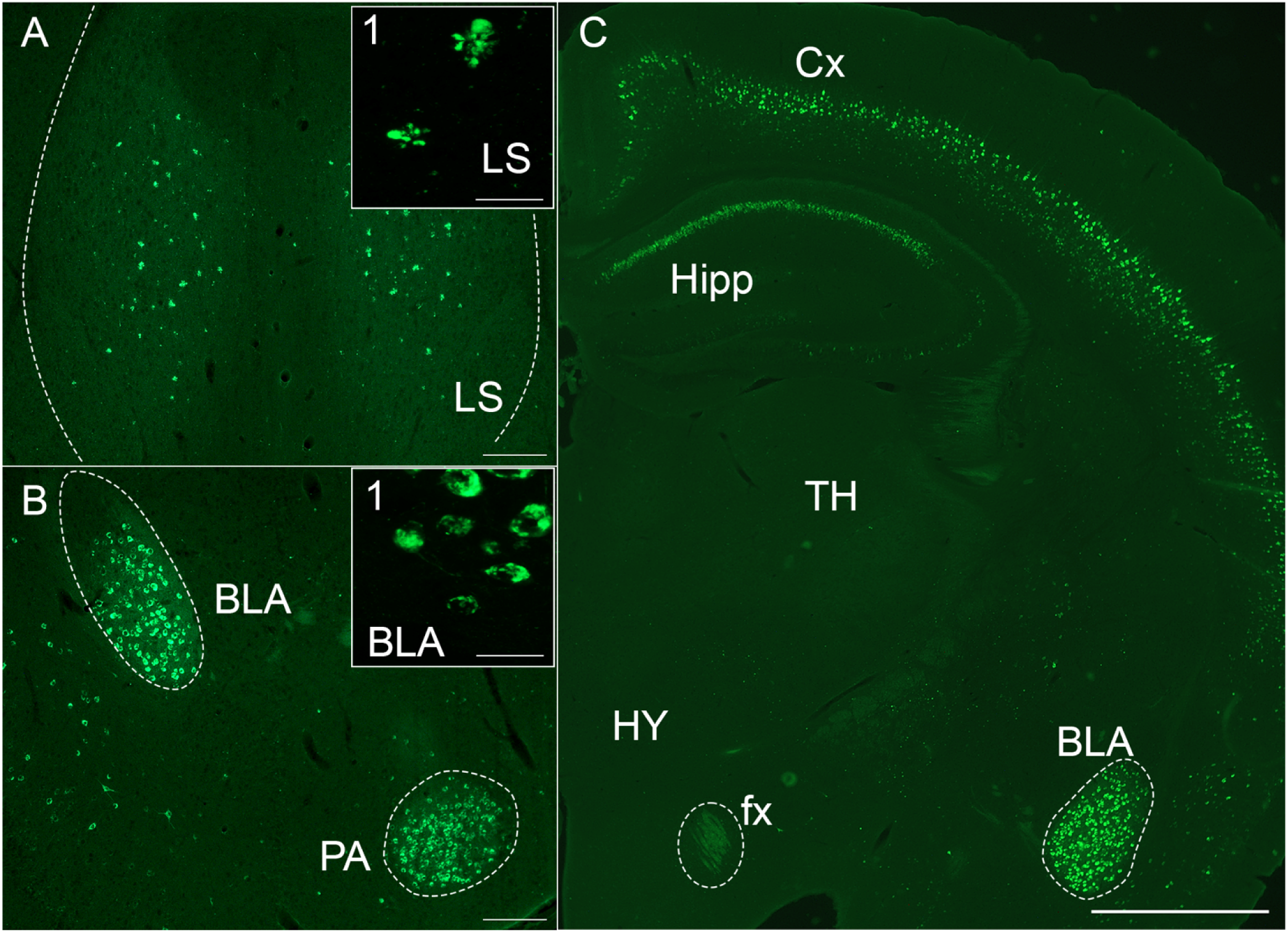
Subcortical structures exhibiting hAPP immunofluorescence in 2-month-old 5xFAD^+/−^/Rosa^tdT^ mice. **(A)** Bilateral fluorescence in the lateral septum. **(A1)** Higher magnification image of rosette shaped 1D1 fluorescent particles. **(B)** Immunofluorescence of hAPP within the basomedial amygdala (BLA) and posterior amygdala (PA) **(B1)** Higher magnification view of 1D1 immunofluorescence in the BLA. **(C)** Immunofluorescence of hAPP in the cortex (Cx), hippocampus (Hipp), BLA and in columns of the fornix. No apparent hAPP immunofluorescence in the thalamus (TH) and hypothalamus (HY). **(A1 & B1)** Imaged on Zeiss LSM 900 Airyscan 2. Scale bar: 200µm in **A&B**; 30µm in **A1 & B1**; 1mm in **C**.

As expected, based on previously reported patterns of Thy1-driven expression, there were only scattered populations of 1D1-labeled neurons in most subcortical regions including thalamus, hypothalamus and brainstem (**Fig. 3C**). Of note, however, some neurons in the red nucleus did exhibit strong immunofluorescence for hAPP along with other scattered populations of neurons in the brainstem and spinal cord. **Fig. 4A** illustrates a section through the midbrain at the level of the superior colliculus and red nucleus.

**Figure 4.**
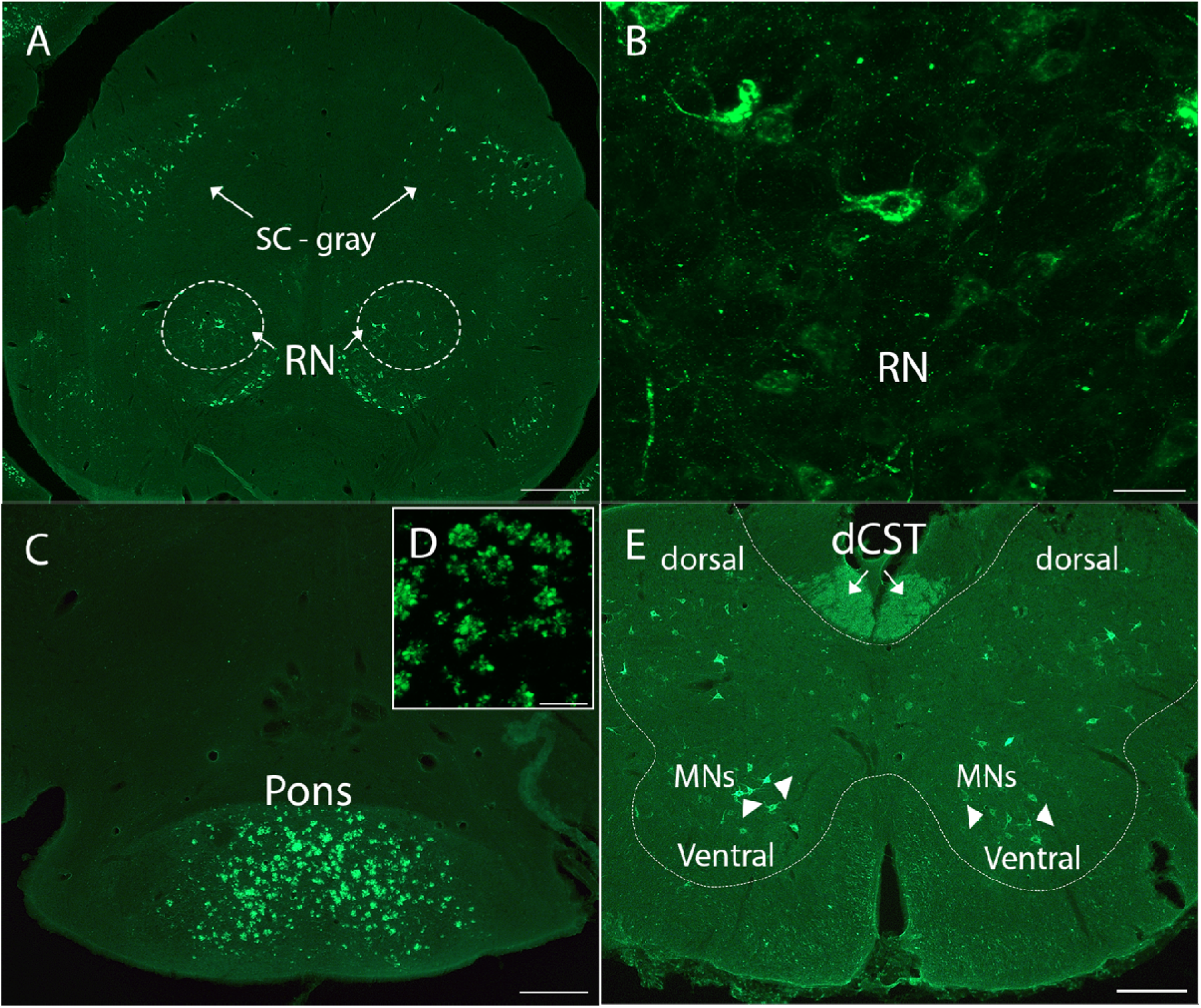
Human APP immunofluorescence in the brain stem and spinal cord of a 2-month-old 5xFAD^+/−^/Rosa^tdT^ mouse. **(A)** hAPP expression in the red nucleus (RN) and in the gray matter of the super colliculus (SC – gray). **(B)** Higher magnification of the RN. **(C & D)** 1D1 immunofluorescence in the pons, seemingly extracellular. **(E)** hAPP labeling of axons in the descending corticospinal tract in the dorsal column of the spinal cord, denoted by the arrows. Sparse labeling of alpha-motor neurons (MNs) in the ventral horn of the s cord, denoted by arrowheads. Scale bar: 200µm **in A, C & E**; 30µm in **B & D.**

1D1-positive particles were particularly prominent in the pons (**Fig. 4C&D).** Neurons in the pontine nuclei project to the cerebellum via the middle cerebellar peduncle where they terminate as mossy fibers on cerebellar granule neurons. The middle cerebellar peduncle is one of the many white matter tracts affected in humans with AD (Toniolo et al., 2020). Although there are many neuronal cell bodies within the pons (pontine nuclei), 1D1 fluorescence appears to be extracellular **(Fig. 4D)**, as in the lateral septum (**Fig. 3A**). In considering the source of these extracellular particles, it should be noted that CST axons terminate on neurons in the pons and pass through the pons en route to the spinal cord. These CST axons originate from 1D1-positive neurons in layer V of the cortex.

In the cervical spinal cord, there was sparse labeling of neuronal cell bodies in the gray matter, including a few large motoneurons in the ventral horn (MNs, **Fig. 4E**). Axons in the ventral part of the dorsal column in the position of the descending corticospinal tract were labeled, consistent with their origin from 1D1-positive CSNs in cortical layer V (dCST, **Fig. 4E)**. Notably, there were no 1D1 fluorescent particles that appeared to be extracellular in the spinal cord at 2 months of age (**Fig. 4E**).

### 3.2 Pattern of transgene expression in older 5xFAD^+/−^/Rosa^tdT^ mice

It was of interest to determine how 1D1 labeling patterns would change as neurons begin to degenerate in older 5xFAD mice. For this, we immunostained brain sections from 7-month-old and older 5xFAD^+/−^/Rosa^tdT^ mice (see Table 1). Notably, the labeling in older 5xFAD^+/−^/Rosa^tdT^ had a different appearance from that observed at 2-3 months of age. In addition to 1D1 fluorescence that appeared to be within neuronal cell bodies (**Fig. 5A&C**), there was a dramatic increase in fluorescent particles that appeared to be extracellular, presumably representing extracellular accumulations of 1D1-labeled hAPP (**Fig. 5B, D**). The 1D1-positive particles appear as irregular or rosette-shaped clusters of particles (traced, **Fig. 5D**), as puncta (circled, **Fig. 5G&H)**, or as threads (arrowheads, **Fig. 5G&H**) similar to Aβ positive threads reported in 5xFAD mice (Chu et al., 2017). These results point to an apparent extracellular accumulation of hAPP as mice age. In what follows, we first consider the areas in which neurodegeneration has been previously reported in 5xFAD mice, layer V of the cortex and subiculum, and then other regions.

**Figure 5.**
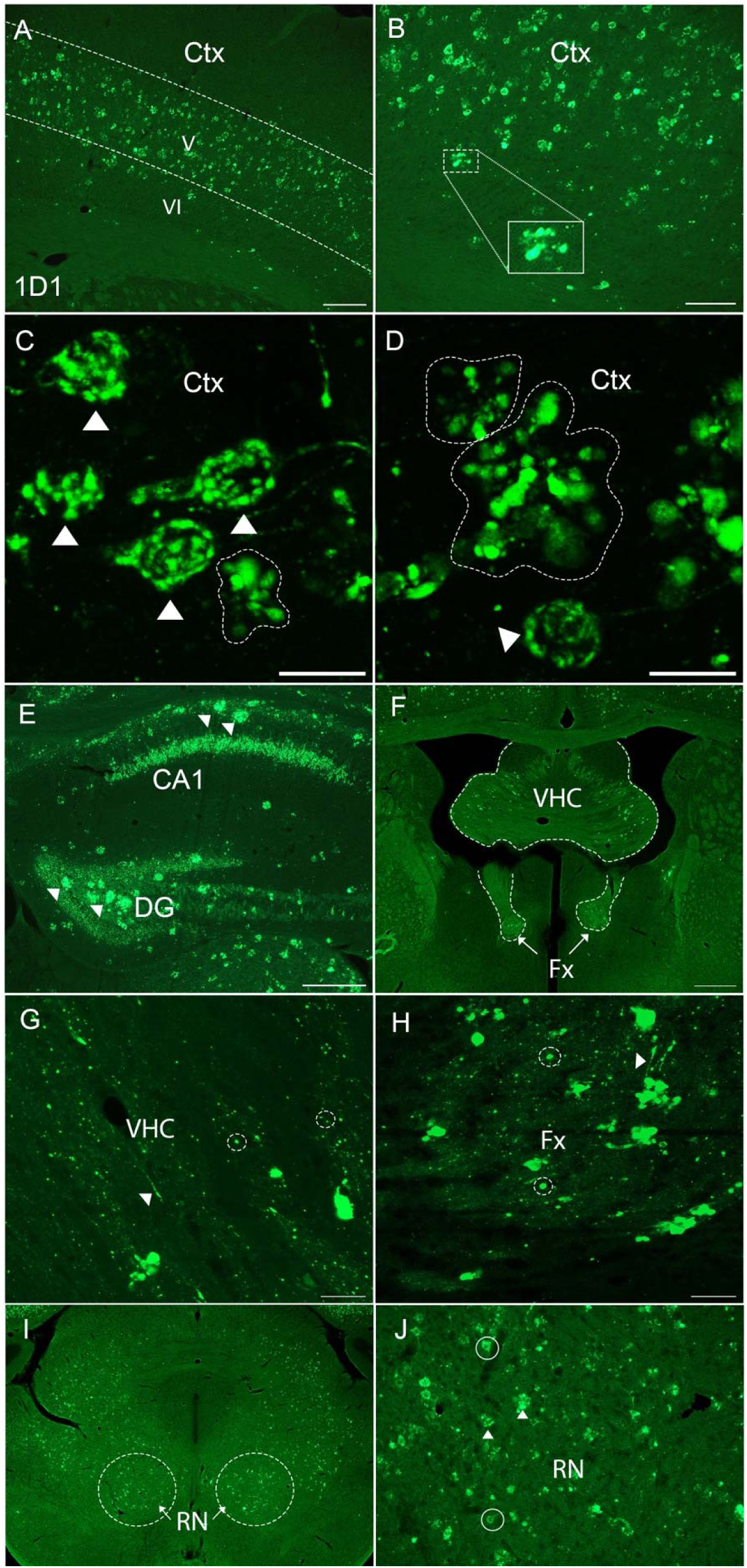
1D1-positive extracellular particles in 7-month-old 5xFAD^+/−^/Rosa^tdT^ mouse. **(A)** 1D1 fluorescence primarily in cortical layer V. **(B)** Higher magnification image of 1D1 immunofluorescence in neurons and rosette-shaped clusters within the primary motor cortex. **(C & D)** 1D1 immunofluorescence throughout neurons in the cortex (arrowheads) and 1D1 positive rosette-shaped clusters in the cortex (traced). **(E)** 1D1 immunofluorescent rosette shaped clusters in CA1 and dentate gyrus (arrowheads). **(F)** View of 1D1 immunofluorescence in the ventral hippocampal commissure (VHC) and the descending column of the fornix (Fx) showing fluorescent fibrils (arrowheads, **G & H**) along with immunofluorescent puncta (circled, **G & H**). **(I & J)** Fluorescence in the red nucleus (RN) appears to be in neurons (**J**, denoted by the circles) and extra-neuronal (**J**, denoted by the arrowheads. Images in panels C and D were taken on the Olympus FV3000 Scanning Confocal Microscope and images in panels G and H were taken on the Zeiss LSM 900 Airyscan 2. Scale bar: 200µm in **A**; 100µm in **B**; 20µm in **C & D**; 300µm in **E**; 500 µm in **F**; 30µm in **G & H**; 500µm in **I**; and 100µm **J**.

In the sensorimotor cortex, primary motor cortex (M1) and primary somatosensory area (S1), 1D1-positive neurons were still evident in layer V in older mice (arrowheads, **Fig. 5C&D**). Overall, however, there were a larger number of extracellular clusters in both layer V (encircled, **Figure 5A**) and in deeper cortical layers (see **Figure 5B** inset). Figure 5D illustrates a confocal image of clusters in layer V. The size of the clusters ranges up to nearly 100µm, larger than the size of a neuronal cell body. Importantly, clusters of 1D1 fluorescent particles below layer V were not present in young mice (**Fig. 1D&F**), indicating progressive accumulation of extracellular hAPP as the mice age. Of relevance, extracellular amyloid-β plaque density has also been reported to progress with age in the cortex of 5xFAD mice (Oblak et al., 2021).

In the CA1 region of the hippocampus, 1D1 immunofluorescence was both intraneuronal and extracellular, with extracellular accumulations being more prominent at 10 months (**Fig. 5E**, coronal section from 7-month-old mouse; **Fig 2B, C & D**, horizontal sections from 10-month-old mouse). Although, there was little to no hAPP fluorescence in the granule cell layer of the dentate gyrus of 2-3-month-old mice (**Fig. 1E**; **Fig. 2A**), there was moderate hAPP IF in the granule cell layer in 7-month-old mice that appeared to be intracellular (**Fig. 5E**). This may coincide with the reported increase in mouse and human RNA transcripts of APP as 5xFAD mice age (Forner et al., 2021). There were also a few large extracellular 1D1-positive particles in the hilus of the dentate gyrus in older mice (arrowhead in dentate gyrus, **Fig. 5E).** These findings are consistent with a recent study involving immunostaining using a different antibody against human and mouse APP (Anti-APP A4 Antibody, a.a. 66-81 of APP (NT), clone 22C11) reporting no intracellular APP in dentate granule cells at 12 weeks of age in 5xFAD, although extracellular plaques were present at that age (Mabrouk et al., 2023).

The pattern of 1D1 IF in the entorhinal cortex of older mice was distinctly different than in young mice. Although there were still almost no 1D1-positive neuronal cell bodies in superficial layers (I-V), there was a dramatic increase in 1D1-IF staining in deep layers adjacent to the white matter, especially in the lateral entorhinal cortex (**Fig. 2C**). Two noteworthy points: 1) The 1D1-positive elements in the deep layers appeared to be primarily extracellular, instead of inside cell bodies (arrowhead in layers V/VI, **Fig. 2C**); 2) The deeper layers are the site of termination of axons from the hippocampus and subiculum. The superficial layers also had a few scattered 1D1-positive clumps (arrowhead in layers III/IV, **Fig. 2C**).

In subcortical areas of older mice, fluorescently labeled particles were present in the red nucleus, midbrain reticular nucleus and superior colliculus, often in rosette-shaped clusters. These appeared to be extracellular (**Fig. 5I&J**), rather than within neuronal bodies as was the case at the 2–3-month timepoint (**Fig. 4A**). Of note, in the older mice, there appeared to be few 1D1-positive neuronal cell bodies in the red nucleus (more on this below).

A key difference between older vs. young mice were the evident clusters of 1D1-positive particles in several white matter tracts. Examples include the corticospinal tract, especially as it passes through the pons (**Fig. 4C&D**), ventral hippocampal commissure (**Fig. 5F&G**) and the descending column of the fornix (**Fig. 5F&H**). All these tracts contain axons from 1D1-expressing neurons (layer V cortical neurons in the case of the CST, hippocampal and subicular neurons in the case of the ventral hippocampal commissure and fornix). The clusters of 1D1-positive particles in the pons are especially noteworthy because some had a petal-like form **(Fig. 4C&D).** Such petal-like structures have previously been associated with degenerating neuronal cell bodies, but 1D1-positive neuronal cell bodies were not seen in the pons at any age. This is strong presumptive evidence that extracellular 1D1-positive particles and petal-shaped clusters can originate from axons and/or synaptic terminals from hAPP-expressing neurons in distant sites.

Taken together, our results indicate that extracellular clusters of 1D1-positive particles are present in young mice in a few areas and increase in abundance in three general sites as mice age: 1) neuropil regions in areas that contain neurons that express hAPP at high levels; 2) white matter tracts that contain axons from neurons that express hAPP at relatively high levels; 3) areas that receive synaptic connections from neurons that express hAPP at high levels – the pons and the deep layer of the entorhinal cortex.

### 3.3 Tracking neurodegeneration of layer V neurons labeled via retrograde transduction with AAV-rg/Cre

Previous studies have reported age-dependent loss of large pyramidal neurons in cortical layer V and neurons in the subiculum in 5xFAD mice (Oakley et al., 2008). However, these reports on neurodegeneration were based on routine neurohistological stains, such as Nissl stains, that are not specific to neuron type. To accurately track degeneration of a particular neuronal type over time requires that the neurons be selectively labeled, ideally with a reporter that is permanent. For this, we used a paradigm that we have previously deployed to transduce neurons of the corticospinal tract to enable regeneration after spinal cord injury (Metcalfe et al., 2022). The paradigm uses a novel variant of AAV that transports cargo retrogradely, (AAV-retrograde, or AAV-rg). AAV-rg expressing Cre (AAV-rg/Cre) is injected into cervical level 5 (C5) of Rosa^tdT^ reporter mice resulting in selective transduction inducing persistent expression of tdT by neurons in cortical layer V that project to the spinal cord (CSNs) as well as other neuron types that project to the spinal cord.

CSNs in motor and somatosensory cortices have axons that descend through the internal capsule, medullary pyramid, pyramidal decussation, and project to the spinal cord. Therefore, injections of AAV-rg/Cre at C5 result in retrograde transduction and permanent labeling of CSNs in the primary motor cortex (M1), secondary motor cortex (M2) and in a region of the secondary somatosensory cortex (S2). The injection site in the spinal cord and distribution of labeled neurons in the cortex can be conveniently visualized before sectioning in intact brains and spinal cords when viewed via epi-fluorescence imaging with excitation light of approximately 550 nm (**Fig. 6A**). In coronal sections through the sensorimotor cortex, tdT labeling occurs throughout the cell bodies, dendritic arbors and axons of the CSNs (**Fig. 6B&C**), which can be visualized with native tdT fluorescence without immunostaining or can be amplified by immunostaining using antibodies for Red Fluorescent Protein (RFP). The injections of AAV-rg/Cre were done unilaterally into the left spinal cord, resulting in tdT labeling of neurons primarily in the contralateral (right) cortical hemisphere. Unilateral injections were done to have an intra-animal control for some of the histological analyses to be used (below). In some mice, neurons in the ipsilateral cortex were also sparsely labeled, likely due to some spread of AAV-rg/Cre across the midline at the injection site (**Fig. 6B**). The injections resulted in labeling of CSNs within the motor cortex, as seen by the counts along the rostro-caudal axis (**Fig. 6D**). These quantifications were done in the same manner as explained in the methods section and counts were plotted by section, per mouse.

**Figure 6.**
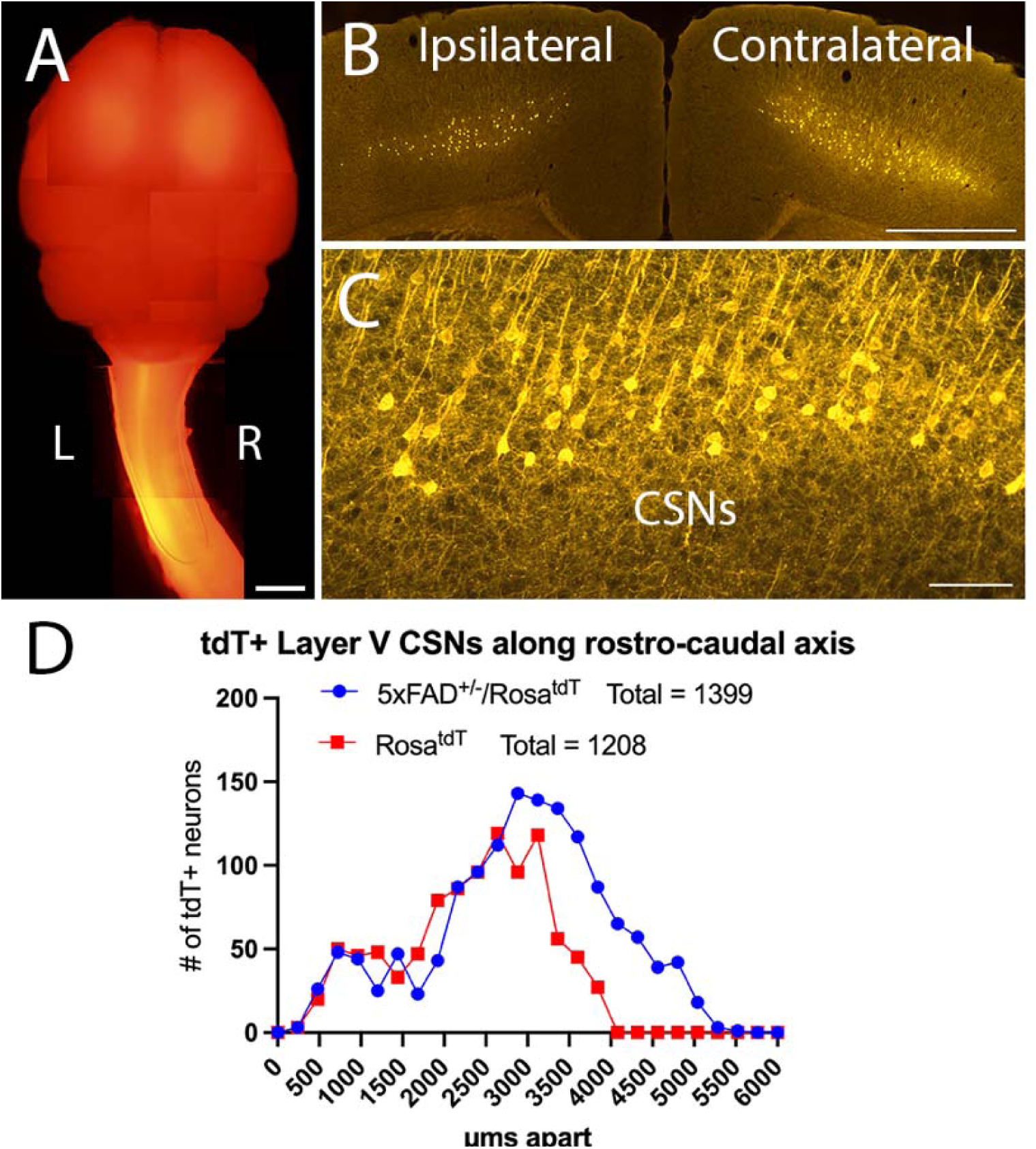
AAV-rg/Cre injections at C5 result in tdT expression in CSNs. **(A)** Epifluorescence of intact Rosa^tdT^ brain and spinal cord. Unilateral injection on left (L) side of the spinal cord resulting in fluorescence primarily on the right (R) cortex. **(B)** Labeling of CSNs, by Red Fluorescent Protein (RFP) IF, primarily on the contralateral side of the cortex and some labeling on ipsilateral side relative to the injection in a Rosa^tdT^ mouse at 2 months of age. **(C)** Precise and substantial labeling of the neuronal bodies and dendritic arbors in Rosa^tdT^ reporter mice, shown by RFP IF. **(D)** Quantification of retrogradely labeled CSNs in 2-month-old 5xFAD^+/−^/Rosa^tdT^ and Rosa^tdT^ mice tdT labeled neurons along the rostro-caudal axis. The number of tdT+ CSNs in 5xFAD^+/−^/Rosa^tdT^ mice are comparable to control Rosa^tdT^ mice. Scale bar: 2mm in **A**; 1mm in **B;** 100µm in **C.**

Neurons that give rise to other spinal pathways are also transduced and express tdT including neurons in the red nucleus, reticular formation, and hypothalamus (Steward et al., 2021). Permanent labeling of cell bodies, axons and dendrites provides a way to track neurodegeneration across age. We leveraged this innovative AAV-mediated delivery tool to define characteristics and timing of neurodegeneration of layer V neurons.

### 3.4 Quantitative consistency in retrograde labeling

To quantify age-dependent CSN loss, it’s important to establish a baseline for the number of transduced CSNs early in life, prior to onset of neurodegeneration. For this, 5xFAD^+/−^/Rosa^tdT^ and control Rosa^tdT^ mice were injected with AAV-rg/Cre between 2-3 months of age and brains were collected 2-3 weeks post-injection, allowing time for tdT to be expressed in transduced neurons. **Fig. 7A** shows a coronal section of a 5xFAD^+/−^/Rosa^tdT^ mouse at 2 months of age. Qualitatively, CSNs appeared morphologically normal in 5xFAD^+/−^/Rosa^tdT^ mice at 2 months (**Fig. 7A**). This 2–3-month timepoint was chosen due to the reported absence of neurodegeneration in 5xFAD mice at this age, allowing accurate quantification the number of transduced neurons at baseline in young 5xFAD^+/−^/Rosa^tdT^ mice.

**Figure 7.**
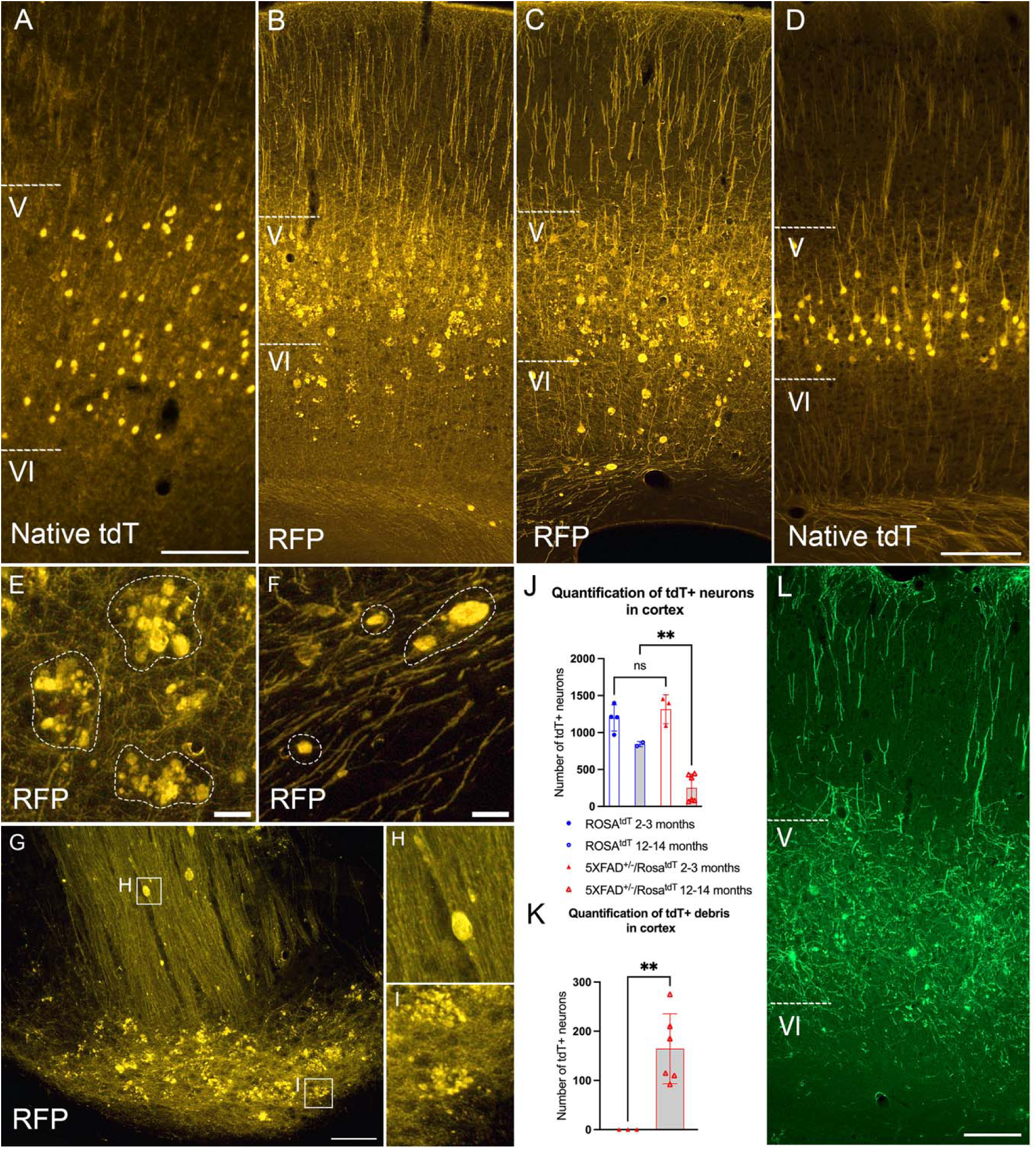
Neurodegenerative changes are labeled by permanent fluorescence. **(A)** CSNs labeled by native tdT in 5xFAD^+/−^/Rosa^tdT^ mouse at 2 months of age **(B)** Expression of tdT, detected by immunofluorescence with an RFP antibody, in primary motor cortex of a 5xFAD^+/−^/Rosa^tdT^ mouse at 8 months of age and **(C)** at 12 months of age. **(D)** No signs of neurodegeneration in the Rosa^tdT^ mouse even at 12 months of age, 10-months post-AAV-rg/Cre injection, shown by native tdT. **(E)** Higher magnification view of the tdT accumulations seen in the cortex and **(F)** dystrophic axons of the tdT positive CSNs in a 5xFAD^+/−^/Rosa^tdT^ mouse at 12 months of age. **(G)** Expression of tdT in the pons of a 14-month-old 5xFAD^+/−^/Rosa^tdT^ reveals axonal varicosities **(H)** and rosette shaped tdT-positive accumulations **(I)**, indicating neurodegeneration. **(J)** Quantifications of tdT-positive CSNs in 5xFAD^+/−^/Rosa^tdT^ and Rosa^tdT^ mice at a young (2-3 months) and older age (12-14 months). Mean number of tdT labeled neurons differed significantly among Rosa^tdT^ and 5xFAD^+/−^/Rosa^tdT^ at 12-14 months of age. There was no significant difference between Rosa^tdT^ and 5xFAD^+/−^/Rosa^tdt^ at 2-3 months (one-way ANOVA: F = 34.15, DF (3, 11), p <0.0001; Sidak’s multiple comparisons test: Rosa^tdT^ at 2-3 months vs. 5xFAD^+/−^/Rosa^tdt^ at 2-3 months, DF = 11, p = 0.6423; Rosa^tdT^ at 12-14 months vs. 5xFAD^+/−^/Rosa^tdt^ at 12-14 months, DF = 11, p = 0.0037. Bars represent mean ± SD. **(K)** Age-related accumulation of tdT-positive debris shown by quantification of 5xFAD^+/−^/Rosa^tdT^ mice at a young (2-3 months) and older age (12-14 months). Debris was not present in Rosa^tdT^ mice. Mean number of tdT labeled debris was significantly greater in 5xFAD^+/−^/Rosa^tdT^ at 12-14 months of age compared to 5xFAD^+/−^/Rosa^tdt^ at 2-3 months of age. Bars represent mean ± SD. Unpaired t-test: t = 3.864, DF =7, p = 0.0062. **(L)** AAV-rg/GFP injected into the cervical spinal cord of 5-month-old 5xFAD^+/−^/Rosa^tdT^ and Rosa^tdT^ mice and mice were allowed to survive until 8 months of age. Scale bar: 100µm in **A**; 200µm in **B-D**; 50µm in **E & F**; 100µm in **G**; 200µm in **L.**

Counts of tdT-labeled CSNs were done on a 1-in-8 series of coronal sections spaced 240µm apart encompassing the rostro-caudal extent of the sensorimotor cortex (**Fig. 7J**). For counts in the 2–3-month timepoint, 12 cases were excluded which labeling was sub-optimal determined by fewer retrogradely-transduced neurons than expected based on our previous studies. In the 12–14-month timepoint group, counts were only excluded if there weren’t any labeled tdT neurons which would indicate a faulty injection. Counts of the total number of tdT-labeled neurons in the 1-in-8 series of sections from young mice ranged from 1000 to 1500 (**Fig. 7J**). The percent variability of the data in 5xFAD^+/−^/Rosa^tdT^ mice is approximately 14.88% and in Rosa^tdT^ mice it is approximately 10.47%. This information provides the baseline for age-dependent CSN degeneration.

#### TdT labeling is permanent

To track neurodegeneration across age, it is important that neurons are permanently labeled with tdT. In both control Rosa^tdT^ and 5xFAD^+/−^/Rosa^tdT^ mice, expression of Cre recombinase removes the stop cassette resulting in robust tdT expression. In previous studies, we have observed robust tdT labeling in CSNs up to 1 year after intra-spinal cord injections of AAV-rg/Cre in Rosa^tdT^ mice. We confirmed robust tdT-labeling of CSNs 6 months after AAV-rg/Cre injection, done at 2 months of age, in 5xFAD^+/−^/Rosa^tdT^ (**Fig. 7B**). In the cortex, tdT-labeling was present throughout the cell bodies, dendrites and axons of layer V CSNs. Even after 10 months of expression, at 12 months of age (**Fig. 7C**) tdT labeling was almost entirely restricted to pyramidal neurons in layer V indicating limited, if any, transneuronal transduction of other neuron types or glia in the cortex (**Fig. 7B&C**). Importantly, when AAV-rg/Cre was injected in Rosa^tdT^ at 2 months of age and the brain tissue was collected 10 months of age, CSNs appeared healthy with no signs of degenerative changes (**Fig. 7D**).

### 3.5 Characteristics of age dependent degeneration of layer V CSNs in 5xFAD^+/−^/Rosa^tdT^ mice

Complete and permanent tdT labeling of cell bodies, axons and dendrites following AAV-rg/Cre injections in 2-month-old 5xFAD^+/−^/Rosa^tdT^ mice allowed us to track degeneration that became apparent as the mice aged. At 8 months of age, tdT labeled CSNs exhibited clear dendritic and axonal dystrophy (**Fig. 7B**). In layers III and above, tdT-positive dendrites appeared irregular in form. Malformations of neuronal processes due to degeneration are termed neuritic dystrophies and are accompanied, or followed by, a loss of dendritic spines. In layers VI and below, large bulbous tdT-positive varicosities were evident along tdT-positive axons in the sub-cortical white matter (**Fig. 7B&C, F**). All these signs of neurodegeneration were more prominent at 12 months of age, including increases in axonal varicosities along CSN axons in layer VI and large varicosities of similar appearance that were not directly along a labeled axon in layer VI (**Fig. 7F**, circled). These manifestations of neurodegeneration were accompanied by decreases in the number of tdT-positive neuronal cell bodies in layer V (see below for quantitative analyses).

A conspicuous feature of labeling in older 5xFAD^+/−^/Rosa^tdT^ mice was the presence of rosette-shaped clusters of extracellular accumulations of tdT that resembled the clusters of 1D1-positive particles (**Fig. 7B, C, E**). In the motor cortex, the tdT particles were primarily below the tdT-positive CSNs in cortical layer VI with minimal presence in layer V. Notably, the tdT-labeled particles were found only in areas of the cortex where tdT labeled CSNs were present. The primary location of the particles, along with their increase in quantity coinciding with a loss of tdT-labeled neurons, suggest that the particles derive from axonal and/or dendritic fragments of tdT labeled neurons that are degenerating. Hence, we term the particles “degeneration debris” from degenerating CSNs.

Accumulations of tdT-positive particles were especially prominent in the pons where axons from tdT labeled CSNs traverse (**Fig. 7G**). The tdT-positive particles were similar in appearance similar to the 1D1-positive particles in the pons illustrated in Figure 5 and included rosette-shaped clusters (**Fig. 7G&I).** It’s important to note that tdT labeled neuronal bodies were not observed in the pons at any age. We also observed large axonal varicosities in the tegmentum dorsal to the pons where tdT-labeled descending CST axons were present (**Fig. 7G&H**). The presence of both extracellular tdT-positive accumulations and 1D1 particles in the pons further supports the interpretation that these originate from axons from layer 1S1-expressing layer V CSNs.

#### Quantification of neurodegeneration

Having established that labeling of CSNs is *permanent* and selective, and signs of neurodegeneration were present, we quantified these features of neurodegeneration as mice age. Brain sections were collected from 5xFAD^+/−^/Rosa^tdT^ and Rosa^tdT^ mice between 2-4 and 12-14 months of age, tdT-labeled CSNs were quantified in a one-in-8 series of sections. The counts revealed a decrease in the number of tdT-positive CSNs at 12-14 months of age in 5xFAD^+/−^/Rosa^tdT^ mice (**Fig. 7J**). In contrast, in control Rosa^tdT^ mice, there was no decrease in the number of tdT-labeled layer V CSNs (**Fig. 7J**). These data confirm and extend previous reports of age-dependent degeneration of layer V CSNs in 5xFAD^+/−^/Rosa^tdT^ mice, document that some of these are CSNs, and reveal that degeneration can be quantitatively tracked by tdT labeling.

In parallel with the decline in tdT-labeled layer V CSNs, tdT labeled extracellular particles in the sensorimotor cortex became more prominent as the mice aged. We quantified the particles in cortical layer V and VI of 5xFAD^+/−^/Rosa^tdT^ mice between 2-4 and 12-14 months of age using the analyze particles plug-in on ImageJ. The inclusion criteria accounted for the various sizes and shapes ranging from large clumps to smaller particles. The quantifications demonstrated that tdT particles increased with age in 5xFAD^+/−^/Rosa^tdT^ mice as CSN counts decreased **(Fig. 7K)**.

We were curious whether a different fluorescent protein expressed by CSNs would also accumulate in extracellular particles as the neurons degenerate. For this, CSNs were labeled by injecting AAV-rg/GFP into the cervical spinal cord of 5-month-old 5xFAD^+/−^/Rosa^tdT^ and Rosa^tdT^ mice and mice were allowed to survive until 8 months of age. Immunostaining for GFP revealed GFP-positive layer V pyramidal neurons along with spherical particles that appeared to be dendritic varicosities in layer IV and above and axonal varicosities below layer V and in the subcortical white matter (**Fig. 7L**). However, there were no clusters of GFP particles resembling the rosette-like accumulations of tdT-positive particles below layer V (**Fig. 7B, C, E**). The overall distribution of tdT-labeled debris resembled the distribution of extracellular accumulations of 1D1-positive particles suggesting possible co-localization. To investigate whether the tdT accumulations co-localized with the 5xFAD transgenes, we performed Co-IF for hAPP, using the FluoReporter^™^ FITC, and Red Fluorescent Protein to label tdT. The IHC was done on tissue from 5xFAD^+/−^/Rosa^tdT^ and Rosa^tdT^ mice an *older* age (12 months of age) when both tdT- and 1D1-stained accumulations of particles were prominent. Co-IF revealed that tdT accumulations were often colocalized with extracellular hAPP (**Fig. 9A-C**). This co-localization supports the conclusion that both the hAPP and tdT-positive extracellular particles originate from tdT-labeled layer V CMNs expressing hAPP.

We were also curious to assess whether neurons in the red nucleus, some of which express hAPP (see Fig. 4A&B; Fig. 5I&J above) would show signs of degeneration. Large numbers of tdT labeled neurons were seen in the red nucleus in young mice **(Fig. 8A)** but far fewer were seen in 12–14-month-old 5xFAD^+/−^/Rosa^tdT^ mice (**Fig. 8B** illustrates a 12-month-old mouse). Quantifications of tdT-positive neurons in the red nucleus of 2-month-old and 12-month-old 5xFAD^+/−^/Rosa^tdT^ mice demonstrated a loss of tdT-positive neurons (**Fig. 8C**) accompanied by the appearance of tdT-positive debris (circled, **Fig. 8B**). To our knowledge, degeneration of neurons in the red nucleus of 5xFAD has not previously been reported in 5xFAD mice.

**Figure 8.**
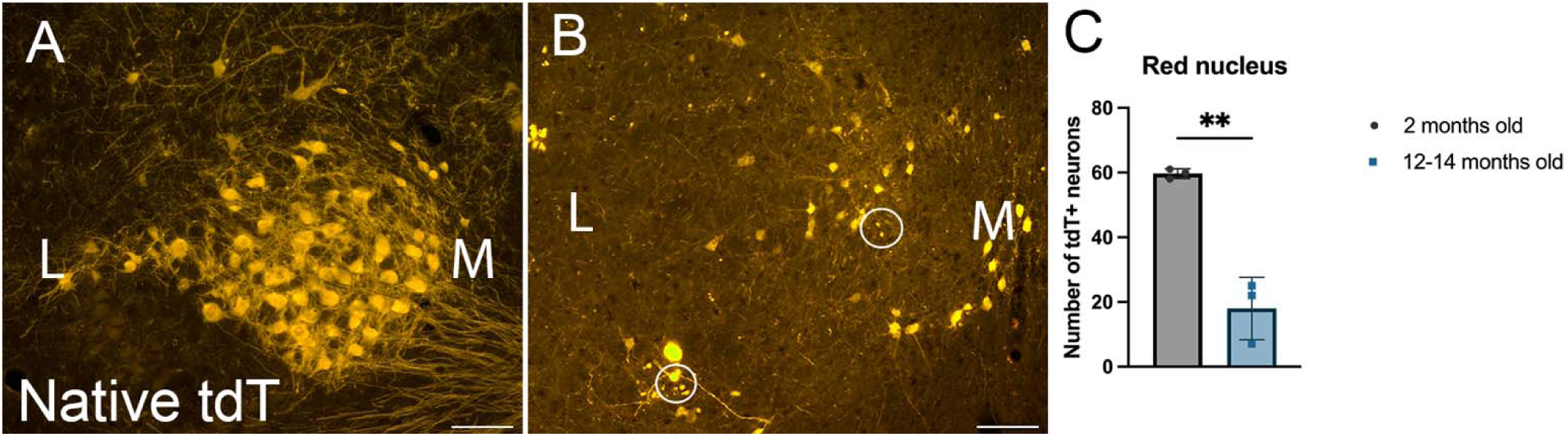
Neurodegeneration shown by fluorescent labeling in 5xFAD^+/−^/Rosa^tdT^. **(A)** Abundant tdT labeled neurons in 2-month-old 5xFAD^+/−^/Rosa^tdT^. **(B)** Fewer tdT labeled neurons in 12-month-old 5xFAD^+/−^/Rosa^tdT^ mouse, along with appearance of debris (circled). Medial (M) and lateral (L). **(C)** Quantifications of neurons in the red nucleus indicate a substantial loss of neurons by 12 months old. Mean number of tdT labeled neurons in the red nucleus was significantly greater in 5xFAD^+/−^/Rosa^tdt^ at 2 months of age than in 5xFAD^+/−^/Rosa^tdt^ at 12-14 months of age. Bars represent mean ± SD. Unpaired t-test: t = 7.391, DF = 4, p = 0.0018. Scale bar: 100µm in **A & B**.

Due to the co-localization of the hAPP transgene and the tdT particles, we explored whether tdT particles were also associated with β-amyloid (Aβ) plaques. For this, sections were co-immunostained for RFP to identify tdT and an antibody that reacts to the N-terminal of Aβ (Amyloid Beta (N3pE) (8E1) Aβ Anti-Human Mouse IgG MoAb) (**Fig. 9D-F**). Additionally, we immunostained for Aβ aggregates with thioflavin-S and observed co-labeling with native tdT fluorescence (**Fig. 9G-I**). As illustrated in Fig. 9D-F, many tdT-positive particles were co-localized with accumulations of Aβ. Taken together, the co-localization studies indicate that at least some of the accumulations of particles contain hAPP, tdT and Aß, suggesting that all three originate from degenerating CSNs. These results also suggest that there must be some unique property of tdT protein that leads to its association with hAPP and Aß in extracellular particles.

**Figure 9.**
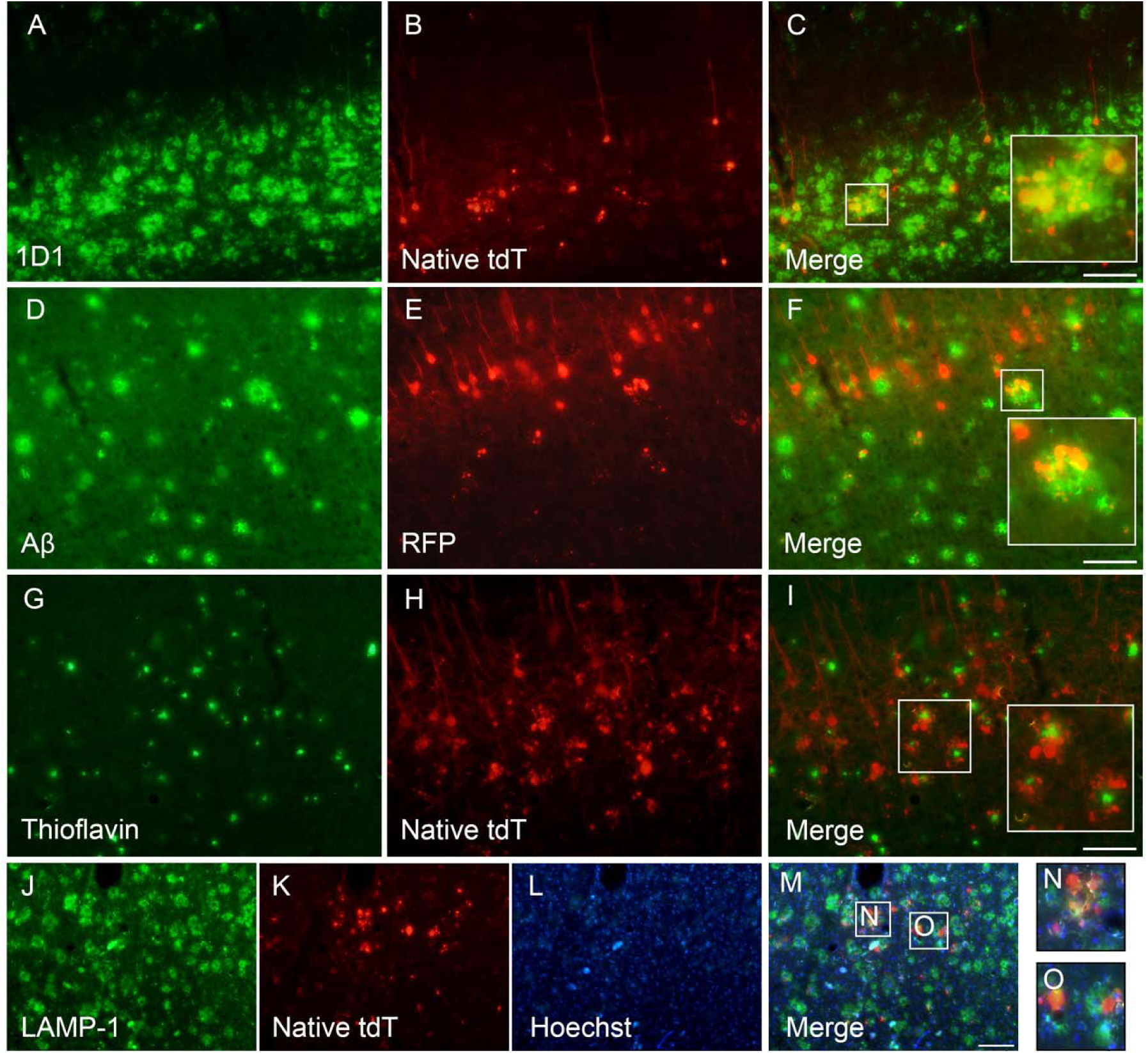
Characterization of tdT accumulations in 12-month-old 5xFAD^+/−^/Rosa^tdT^ mice. **(A-C)** hAPP transgene colocalizes with tdT labled accumulations in cortical layer V/VI. **(D-F)** Aβ appears to colocalize with tdT accumulations in the cortex. **(G-I)** Thioflavin stain shows tdT positive accumulations surrounding amyloid core. **(J-O)** LAMP-1, hoechst and native tdT expression in the cortex. Some tdT accumulations colocalize with Lamp-1 and hoechst **(N)** and others appear isolated, indicating that they are extracellular **(O).** Scale bar: 100µm in **A-O.**

The large size of the tdT-positive clusters (nearly 100 µm in diameter) suggests that they are not inside a cell body, but to assess this more directly, we immunostained sections with Hoechst to label nuclei. As illustrated in Figure 9L-O, the tdT accumulations did not primarily appear to surround a Hoechst-positive nucleus, suggesting that most of the clusters are not inside a cell body. It remains possible that the clusters are within very large axonal swellings described above. In dendritic or axonal swellings, lysosomes have been reported to form clusters (Sharoar et al., 2021). To investigate whether the tdT accumulations are within axonal swellings and associating with lysosomal clusters, we immunostained for LAMP-1 and Hoescht (**Fig. 9J-O**). Some of the smaller tdT-positive accumulations appeared to be associated with Hoechst and LAMP-1 (**Fig. 9M, N**), while the larger tdT clusters did not appear to be associated with either (**Fig. 9M, O**). These data suggest that the accumulations are associated with the hAPP transgene outside of cells and are not within the large axonal varicosities.

It is not surprising that the mutant hAPP is found in the same extracellular particles as Aß, but tdT is an exogenous cytoplasmic protein, so how it gets from the neurons that express the protein into the Aß/hAPP particles is not apparent. There is no compelling reason to predict that tdT, a soluble cytoplasmic protein, would bind to either Aß or hAPP, but this possibility can’t be excluded.

## 4. Discussion

This study provides several novel insights about the nature of age-related neurodegeneration in 5xFAD mice. Key findings are: 1) Location and expression of hAPP transgenes differ between young and old 5xFAD^+/−^/Rosa^tdT^ mice; 2) As 5xFAD^+/−^/Rosa^tdT^ mice age, there is a dramatic increase in extracellular hAPP-positive particles; 3) Extracellular particles containing hAPP are especially prominent in areas containing dendrites, axons and synapses from neurons expressing the hAPP transgenes; 4) Selective retrograde labeling of CSNs reveals extensive degeneration with age, in 5xFAD^+/−^/Rosa^tdT^ mice; 5) Extracellular tdT particles increase with age as tdT+ CSNs decline, providing another quantitative measure to track neurodegeneration of tdT-positive CSNs; 6) There is also neurodegeneration in the red nucleus of 5xFAD^+/−^/Rosa^tdT^ mice, which has not been previously reported on, to our knowledge; 7) As mice age, tdT-positive extracellular particles accumulate in the same areas with extracellular particles containing hAPP; 8) Co-immunostaining reveals substantial co-localization of extracellular hAPP, extracellular tdT protein and Aß-containing plaques. Taken together, our findings establish that some proteins derived from hAPP-expressing neurons accumulate in extracellular particles in areas containing axons and synaptic terminals from hAPP-expressing neurons. The clusters of particles often have a rosette-like or petal shaped structure even in areas that do not contain neuronal cell bodies that express hAPP. In what follows, we discuss implications and caveats of our findings.

### IHC for hAPP reveals previously un-described selectivity in transgene expression

The Thy1-promoter drives transgene expression in 5xFAD mice, and its pattern of expression were largely consistent with previously described patterns of Thy1-YFP expression. However, some aspects of selectivity of expression should be noted: 1) Transgene expression was selective to different neuronal types in the cortex, but varied across cortical areas; 2) Some of the neuronal types that express the transgene at high levels have been reported to exhibit age-related neurodegeneration (neurons in cortical layer V and subiculum) but other neurons that express the transgene at moderately high levels have not been reported to degenerate (neurons in the BLA); 3) There was a notable absence of transgene expression in neurons in the entorhinal cortex that project to the dentate gyrus and hippocampus; 4) 1D1-positive structures were present in some brain regions that appeared to be extracellular rather than intracellular (more below).

### IHC for hAPP reveals previously un-described features of neurodegeneration in 5xFAD mice

The spatial and age-dependent differences in hAPP transgene expression observed in 5xFAD^+/−^/Rosa^tdT^ mice suggests previously un-described, or at least under-described, aspects of neurodegeneration. Expression of hAPP appeared largely intraneuronal in young 5xFAD^+/−^/Rosa^tdT^ mice, except in the lateral septum and pons, but appeared to become primarily extracellular as degeneration occurred in the older 5xFAD^+/−^/Rosa^tdT^ mice. In the older 5xFAD^+/−^/Rosa^tdT^ mice, 1D1-positive particles became prominent in the cortex and subiculum. Notably, the clusters of 1D1 particles in the motor cortex were most prominent in cortical layer VI, below the neurons in layer V that express the transgene at high levels. The 1D1-positive particles also became more abundant in fornix and lateral septum as the 5xFAD^+/−^/Rosa^tdT^ mice aged. The fornix contains axons from the subiculum, where our results reveal high levels of expression of hAPP. The lateral septum receives projections from the hippocampus and the basolateral amygdala, where there is a high level of 1D1 expression in neuronal cell bodies. It’s possible that as 1D1 positive neurons are degenerating they are subsequently releasing hAPP. The presence of APP was previously reported in swollen degenerating axons that were surrounding amyloid plaques and neuritic plaques (Dickson, 1997; DeTure and Dickson, 2019; Mabrouk et al., 2023).

Especially dramatic accumulations of hAPP were observed in the pons. Of note, CST axons from cortical layer V neurons pass through the pons *en route* to the spinal cord and terminate on neurons in the pontine nuclei. However, we did not observe hAPP-labeled neuronal cell bodies in the pons at any age. These findings strongly support the conclusion that extracellular particles of hAPP are deposited by axons and synaptic terminals from hAPP-expressing neurons located in distant structures, not by degenerating hAPP-expressing neuronal cell bodies in the pons.

Some extracellular clusters of hAPP resemble what have previously been termed “PANTHOS”, which were interpreted as the remains of degenerated neuronal cell bodies. The proposed process is that autophagic vacuoles within the neuronal cell body generate a β-amyloid ‘core’ (Lee et al., 2022). If this is the case, then PANTHOS structures should appear only where neuronal cell bodies have degenerated. However, our results reveal rosette-like clusters of 1D1-positive particles (PANTHOS-like) in axon tracts and areas of termination of axons from 1D1-positive neurons in structures that do not contain 1D1-expressing neuronal cell bodies. These results suggest that PANTHOS structures can originate from degenerating axons and synaptic terminals. Our results are consistent with another recent study showing that extracellular Aß develops in the dendritic lamina of the dentate gyrus where the granule cells express the 5xFAD transgene at low levels. These results were interpreted as indicating that extracellular Aß in the dendritic zone of the dentate gyrus originates from axons and terminals of neurons that express the transgene (Mabrouk et al., 2023). Of note, extracellular accumulations of tdT from CSNs also sometimes had a PANTHOS-like appearance (more on this below).

The change in hAPP localization (from intracellular to extracellular) coincides with neurodegeneration of certain neuron types in cortical layer V and the subiculum. Our results characterizing areas of extracellular hAPP deposition may be useful for identifying unreported areas of neurodegeneration in 5xFAD mice.

### Permanent labeling of corticospinal, and other neurons that project to the spinal cord, enabled quantification of neurodegeneration of defined neuron types in 5xFAD mice

We developed the 5xFAD^+/−^/Rosa^tdT^ line to quantify age-dependent neurodegeneration of CSNs and the approach achieved that goal. Counts of tdT-expressing CSNs after retrograde transduction with AAV-rg/Cre revealed loss of about 80.92% of CSNs. Our results also revealed previously undescribed neurodegeneration in the red nucleus, which gives rise to the rubrospinal tract, involved in motor control (Vadhan et al., 2024). The neurons in the red nucleus also receive inputs from the motor cortex and cerebellum. Thus, age-related degeneration of these neurons could contribute to the motor symptoms that develop in 5xFAD mice. Neurodegeneration may be occurring in other areas of the brain where extra-neuronal hAPP immunofluorescence was observed, but our model was created to track neurodegeneration in neurons with projections to the spinal cord. Of note, Cre-dependent induction of tdT provided by 5xFAD^+/−^/Rosa^tdT^ mice could be used for selective labeling of other neuron types in these mice to track neurodegeneration.

### Permanent labeling of corticospinal and other neurons that project to the spinal cord reveals other aspects of neurodegeneration

The 5xFAD^+/−^/Rosa^tdT^ model also revealed previously undescribed extracellular accumulation of some exogenous proteins expressed by neurons undergoing degeneration in 5xFAD mice. It is well-established that APP accumulates in extracellular particles in 5xFAD mice, as well as in human AD where it is associated with Aß plaques. However, we were surprised that tdTomato, expressed by degenerating neurons, also accumulates in extracellular particles and co-localized with hAPP. It is exceedingly unlikely that the extracellular tdT could have come from any source other than the tdT-expressing neurons. We were further astonished that tdT continued to generate native fluorescence in these extracellular locations’ weeks, and perhaps months, after it had been produced by tdT-expressing neurons. It remains unclear how tdT survives as a functional fluorescent protein for this duration of time. Additionally, the presence of petal-like accumulations of extracellular tdT, in white matter tracts containing tdT labeled CST axons, supports the speculation that petal-like structures are the remains of degenerated axonal swellings. This appearance of axonal varicosities in tdT labeled layer V CSNs, in conjunction with high 1D1 labeling in layer V, provides compelling evidence that axonal varicosities are on axons from 1D1-expressing CSNs.

Because of tdT being a soluble protein, it is also unclear how it remains in the same location as extracellular hAPP granules. This suggests there must be a strong interaction between the tdT protein and hAPP. This could be part of the intracellular pathophysiology due to expression of the 5xFAD transgenes. For example, the protein encoded by the transgene may bundle and perhaps cross-link with other proteins. It may be possible to use the association between extracellular tdT and hAPP as a biochemical tool to further characterize the components of the extracellular accumulations.

We also identified accumulations of GFP in the 5xFAD^+/−^/Rosa^tdT^ mice that were injected with AAV-rg/GFP. The accumulations appeared to be within axons of GFP labeled neurons and were not nearly as abundant as the accumulations seen with tdT labeling. The more prominent presence of tdT, when compared to GFP, may be due to its longer half-life and greater stability. Additionally, in our models, tdT is driven by a constitutive promoter whereas GFP is introduced via a viral vector resulting in a reliance of vector stability for its expression.

Our work adds to the reports on the association of degenerating particles with Aβ by demonstrating that hAPP protein is also present in the rosettes or dystrophic neurites containing Aβ. This is relevant because the APP protein can be cleaved by ß-secretases, and subsequent γ-secretases, leading to Aβ formation (Haass et al., 2012). Thus, our results also contribute to the hypothesis that neurons die and convert into ‘extracellular’ plaques (D’Andrea et al., 2001).

### Limitations of 5XFAD as a model of human AD

A limitation of the 5xFAD mouse model is that it is not representative of how AD manifests in humans. First, the transgenes are driven by the Thy-1 promoter, so expression is limited to certain neuron types. In contrast, in humans carrying FAD mutant alleles, most or all neurons express the gene although levels of expression probably vary by neuron type. Second, 5xFAD mice carry a total of 5 different FAD mutant alleles in two different genes, which is certainly not the case in any human. FAD makes up a small amount of the cases of AD, so the 5xFAD mouse is not representative of most human cases. Third, human AD is characterized by a slow progression of pathology that includes extracellular amyloid-β plaques and intracellular neurofibrillary tau tangles (Knopman et al., 2021). The 5xFAD model, an amyloidosis model, exhibits a rapid and abundant amyloid-β burden. While phosphorylated tau aggregates have been reported in 5xFAD mice (Shin et al., 2021), they aren’t a large component of the 5xFAD pathology. Of note, using the same antibody as Shin et al., we did not detect phospho-Tau in our 5xFAD mice (*unpublished data*). Finally, 5xFAD mice display prominent age-dependent motor deficits that are not a prominent feature of AD in humans (Masse et al., 2021).

It is also important to note that our transgenic line of 5xFAD^+/−^/Rosa^tdT^ was generated by crossing different strains and thus has a different genetic background than the commercially available 5xFAD-B6SJL background mice. The 5xFAD^+/−^/Rosa^tdT^ mice and Rosa^tdT^ mice have been propagated in our facility over multiple generations, which results in genetic background differences that may introduce variability in disease pathology and behavior.

For all these reasons, our findings described herein define aspects of neurodegeneration in 5xFAD mice and perhaps even specifically 5xFAD^+/−^/Rosa^tdT^ mice. Some of the features of pathophysiology may not represent the situation in human AD, including FAD. Despite this caveat, our findings do reveal novel aspects of neurodegeneration and new conclusions that likely apply to 5xFAD mice in general until reported otherwise.

## Acknowledgements

Supported by NIH grant NS047718 to OS. Thank you to Ardi Gunawan for skilled mouse surgery.

## Author contributions

OS and DG designed research, DG performed research and analyzed data. DG and OS wrote the paper.

